# Heightened efficacy of anidulafungin when used in combination with manogepix or 5-flucytosine against *Candida auris in vitro*

**DOI:** 10.1101/2022.12.16.520848

**Authors:** Larissa L.H. John, Darren D. Thomson, Tihana Bicanic, Martin Hoenigl, Alistair J.P. Brown, Thomas S. Harrison, Elaine Bignell

**Affiliations:** Medical Research Council Centre for Medical Mycology, School of Biosciences, University of Exeter, Exeter, UK; Institute of Infection and Immunity, St George’s University of London, London, UK; Division of Infectious Diseases, Medical University of Graz, Graz, Austria; BioTechMed, Graz, Austria; Translational Medical Mycology Research Unit, ECMM Excellence Center for Medical Mycology, Medical University of Graz, Graz, Austria; Clinical Academic Group in Infection and Immunity, St George’s University Hospitals NHS Foundation Trust, London, UK

**Keywords:** *Candida auris*, antifungal combination, anidulafungin, flucytosine, manogepix, synergy

## Abstract

*Candida auris* is an emerging, multi-drug resistant fungal pathogen that causes refractory colonisation and life-threatening invasive nosocomial infections. The high proportion of *C. auris* isolates that display antifungal resistance severely limits treatment options. Combination therapies provide a possible strategy to enhance antifungal efficacy and prevent the emergence of further resistance. Therefore, we examined drug combinations using antifungals that are already in clinical use or undergoing clinical trials. Using checkerboard assays we screened combinations of 5-flucytosine and manogepix (the active form of the novel antifungal drug fosmanogepix) with anidulafungin, amphotericin B or voriconazole against drug resistant and susceptible *C. auris* isolates from clades I and III. Fractional inhibitory concentration indices (FICI values) of 0.28-0.75 and 0.36-1.02 were observed for combinations of anidulafungin with manogepix or 5-flucytosine, respectively, indicating synergistic activity. The high potency of these anidulafungin combinations was confirmed using live-cell microfluidics-assisted imaging of fungal growth. In summary, combinations of anidulafungin with manogepix or 5-flucytosine show great potential against both resistant and susceptible *C. auris* isolates.

## Introduction

*Candida auris* is an emerging fungal pathogen that causes nosocomial invasive infections and that is difficult to eradicate following colonisation of hospitalised patients (1). *C auris* was first identified in 2009 in Japan, but since then outbreaks have been observed on most continents (1, 2). *C. auris* strains have been subdivided into four genetic clades, the South Asian (I), East Asian (II), South African (III) and South American (IV) clades (3), with a potential fifth Iranian clade identified more recently (4). The organism colonises the skin and can lead to mucosal or bloodstream infections, predominately in immunocompromised hosts (1). Invasive *C. auris* infections are associated with mortality rates between 28% and 60%, and treatment failure due to antifungal resistance is often observed (1, 3, 5–11).

To date, only four classes of antifungal drug are available for the treatment of invasive fungal infections: the azoles, polyenes, echinocandins and the nucleoside analogue 5-flucytosine. 5-flucytosine has high oral bioavailability with high activity against *C. auris*, but it is not generally used in monotherapy due to the rapid emergence of resistance (12). Current guidelines recommend echinocandin treatment as first line therapy for invasive candidiasis and for *C. auris* infection in particular (13, 14). However, echinocandin resistance can develop during treatment (15, 16). Resistance to all four existing classes of antifungal has been reported in *C. auris*, with varying drug susceptibilities and resistance mechanisms between clades (17). Around 90 % of *C auris* isolates show resistance to fluconazole with varying susceptibilities to other azoles (3, 6, 9, 18). Resistance to amphotericin B and the echinocandins appears to be less common, having been reported in 13-35 % and 2-7 % of tested isolates, respectively (3, 9, 18). Alarmingly, between 3 % and 41 % of isolates exhibit resistance to two or more antifungal classes (3, 18). Consequently, the Centers for Disease Control and Prevention (CDC) recently added *C. auris* to its list of urgent antibiotic resistance threats (19) and the World Health Organisation (WHO) declared it a critical threat in its fungal priority pathogens list (14).

The limited number of antifungal drugs as well as the increased threat of antifungal resistance in *C. auris* means that novel treatment strategies are urgently needed. Combinations of antifungals with different mechanisms of action provide one proposed therapeutic strategy. Previous *in vitro* studies investigated combinations of echinocandins with azoles or the polyene amphotericin B (20–24) and combinations of 5-flucytosine with the other three antifungal classes in *C. auris* (25–27). These studies observed either synergy or indifference and no antagonism for all of the tested combinations, with variability between *C. auris* isolates. The most promising combinations were azoles combined with echinocandins which, in two studies, resulted in synergy against all tested isolates (20, 23).

Combinations with 5-flucytosine are of particular interest as its combinations with amphotericin B and fluconazole have been shown to be superior to monotherapy in phase III clinical trials against cryptococcal meningitis (28). As a result of these trials, 5-flucytosine is now more widely available globally, including in countries such as South Africa which suffers a high burden of *C. auris* candidemia (28, 29). Echinocandin combinations with 5-flucytosine have been reported to be indifferent in most cases, but these combinations have shown 100% growth inhibition and fungicidal activity against multidrug-resistant isolates (25–27).

None of these studies included the new antifungal fosmanogepix, which has recently completed phase 1 and 2 clinical trials, and is one of several new antifungals in the pipeline that may exhibit activity also against *C. auris* (30). Fosmanogepix is a prodrug that is converted to the active compound manogepix by systemic phosphatases (31). Manogepix inhibits a novel antifungal target, Gwt1, which is involved in the GPI-anchor biosynthetic pathway, leading to a decrease in cell wall-anchored mannoproteins (31). In the present study, we examined combinations of manogepix or 5-flucytosine with anidulafungin, amphotericin B or voriconazole against a range of resistant and susceptible *C. auris* isolates *in vitro*.

## Material and Methods

### Fungal isolates

Twenty-five clinical *C. auris* isolates belonging to clades I, III and IV isolated from 6 patients from a range of sites (blood, urine, respiratory tract, skin) were obtained from the CDC (Table 1). Clade designations were based on whole genome sequencing (Gifford *et al*., in preparation). Isolates were maintained at - 80 °C in 25 % glycerol broth and subcultured on Sabouraud dextrose agar (SDA) at 37 °C for up to 48 h.

**Table 1.**
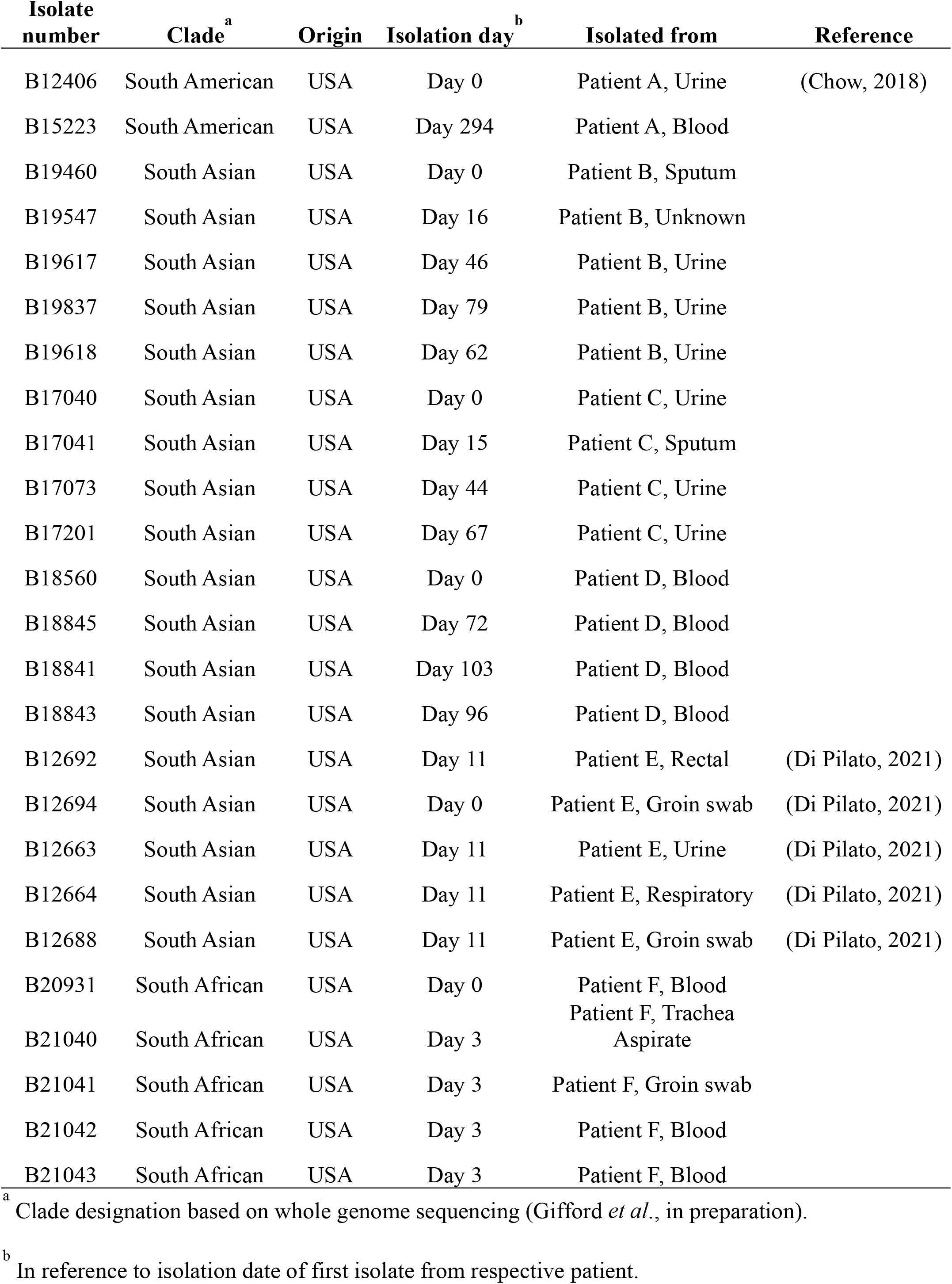
*Candida auris* isolates. (46, 47)

### Antifungal susceptibility testing

Antifungal susceptibility testing was performed using the broth microdilution method according to EUCAST guidelines (32). Anidulafungin (MedChem Express), amphotericin B (Merck), fluconazole (Thermo Scientific), 5-flucytosine (Thermo Scientific), fosmanogepix (MedChem Express), manogepix (MedChem Express) and voriconazole (Sigma Aldrich) were dissolved in 100 % dimethyl sulfoxide (DMSO). The range of antifungal concentrations tested were 0.016 to 8 mg/L for anidulafungin, 0.03 to 16 mg/L for amphotericin B and voriconazole, 0.25 to 128 mg/L for fluconazole, 0.008 to 4 mg/L for 5-flucytosine, 0.004 to 2 mg/L for fosmanogepix and 0.002 to 1 mg/L for manogepix. The minimum inhibitory concentration (MIC) endpoint for amphotericin B was defined as the lowest concentration leading to 90 % reduction in growth compared to the drug-free control (MIC_90_), while MIC_50_ endpoints, measuring 50 % reduction, were used for all other antifungal agents. Tentative CDC breakpoints for *C. auris* were used to define resistance to anidulafungin (≥4 mg/L), amphotericin B (≥2 mg/L), fluconazole (≥32 mg/L) and voriconazole (≥2 mg/L) (https://www.cdc.gov/fungal/candida-auris/c-auris-antifungal.html). A known issue for broth microdilution susceptibility testing of amphotericin B in RPMI medium is the clustering of MICs around the breakpoint of 2 mg/L making it difficult to distinguish resistant and susceptible isolates (33). There are no breakpoints available for 5-flucytosine and fosmanogepix. *Candida krusei* ATCC 6258 and *Candida parapsilosis* ATCC 22019 were used as quality control strains as recommended by the EUCAST guidelines (32). All experiments were performed in triplicate.

### Antifungal combination testing

Interactions of antifungal drugs were tested using checkerboard assays based on EUCAST guidelines (32). The range of antifungal concentrations tested was dependent on the MIC of each isolate, with the highest concentration at 4 x MIC. Columns 3 to 12 of a 96-well microtiter plate were filled with 50 μl of drug A and rows B to H were filled with 50 μl of drug B. Column 1 served as drug-free growth and sterility control. The inoculum was prepared by suspending five distinct colonies from 40-48h-old cultures in distilled water, counting the cell number using a haemocytometer and adjusting inocula to 5 × 10^5^ cells/ml. The plates were inoculated with 100 μl and incubated at 37 °C for 24 h. OD readings were taken after 24 h using a spectrophotometer at 530 nm. All experiments were performed in triplicate.

Two different approaches were applied in the analysis of drug interactions. The fractional inhibitory concentration index (FICI) was calculated as follows:

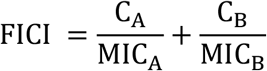

C_A_ and C_B_ are the concentrations of the drugs A and B in combination and MIC_A_ and MIC_B_ are the MICs of the drugs alone. MIC values were rounded to the next highest two-fold concentration if the endpoint was not reached within the tested concentration range. The interaction was considered synergistic for FICI ≤0.5, partially synergistic between >0.5 and <1.0, additive at 1.0, indifferent between >1.0 and <4 and antagonistic >4 (24). In the following, the term “any synergy” refers to FICI values of <1, thereby including complete and partial synergy. In the presence of antagonism, the maximum median FICI values were reported, otherwise minimum median FICI values were given. Additionally, drug interactions were visualised using a response surface analysis approach with Combenefit software (version 2.021) under application of the Bliss independence model (34).

### Microfluidics imaging

*C. auris* B12663 cells were grown and prepared as described above. Inocula were adjusted to 2 × 10^5^ cells/ml. Antifungal mono- and combination treatments were prepared in RPMI 1640 supplemented with glucose to 2 % and 0.165 mol/L 3-(N-morpholino) propanesulfonic acid (MOPS) (RPMI 2%G) at the MIC. CellASIC® ONIX Y04C microfluidic plates (Millipore Merck) were washed with RPMI 2%G by applying 5 psi perfusion for 5 min using the CellASIC® ONIX2 microfluidic system (version 1.0.4 Millipore Merck). Yeasts were loaded into the CellASIC culture chambers by applying 8 psi for 5 s twice (Thomson *et al*., in preparation). Adhered cells were then perfused with RPMI 2%G for 4 h at 1 psi. After 4 h, cells were exposed to the antifungal(s), or to RPMI 2%G for the drug-free control, by applying 5 psi for 5 min, followed by perfusion at 1 psi for 20 h at 37 °C, during which the microfluidic plates were subjected to multi-point 4D imaging on an inverted AxioObserver Z1 microscope (Carl Zeiss). Differential interference contrast (DIC) images were captured with a 20x/0.8NA PlanApochromatic DIC objective and a 16-bit ORCA-Fusion sCMOS camera (Hamamatsu). The area of colonies over time was measured in FIJI 1.53t (35) using an adapted method for migration analysis from Venter and Niesler (36). Briefly, during the time series, colony edges were found (Process → Find Edges), the image blurred fifteen times (Process → Smooth) and inverted (Edit → Invert) before thresholding (Image → Adjust → Threshold: Default) to quantify the total fungal area (Analyse → Analyse Particles). Increases in colony area were used to calculate the doubling times.

## Results

### Antifungal activity against C. auris isolates

The antifungal susceptibility profiles of 25 *C. auris* isolates were determined in order to select a subset of isolates with different drug susceptibilities for antifungal combination testing. The ranges of MIC values for the *C. auris* isolates against the tested antifungals are summarised in Table 2 and Table S1. MIC values for amphotericin B clustered around the breakpoint of 2 mg/L which is a known problem for broth microdilution susceptibility testing of amphotericin B in RPMI medium, making it difficult to distinguish resistant and susceptible isolates (33). Fluconazole showed a large percentage of resistant *C. auris* isolates (96 %; breakpoint ≥32 mg/L) with high MIC values ranging from 4 to ≥128 mg/L, while the other triazole tested (voriconazole) displayed more potent antifungal activity with MICs ranging from 0.06 to 16 mg/L and 40 % resistant isolates (breakpoint ≥2 mg/L). Of all the antifungals tested with an available breakpoint, anidulafungin produced the lowest percentage of resistant isolates (32 %; ≥4 mg/L). The most potent antifungal activity against *C. auris* was observed for manogepix (MIC_50_/MIC_90_, 0.008/0.03 mg/L; range, 0.004-0.03) followed by 5-flucytosine (MIC_50_/MIC_90_, 0.25/0.25 mg/L; range, 0.125-0.25).

**Table 2.**
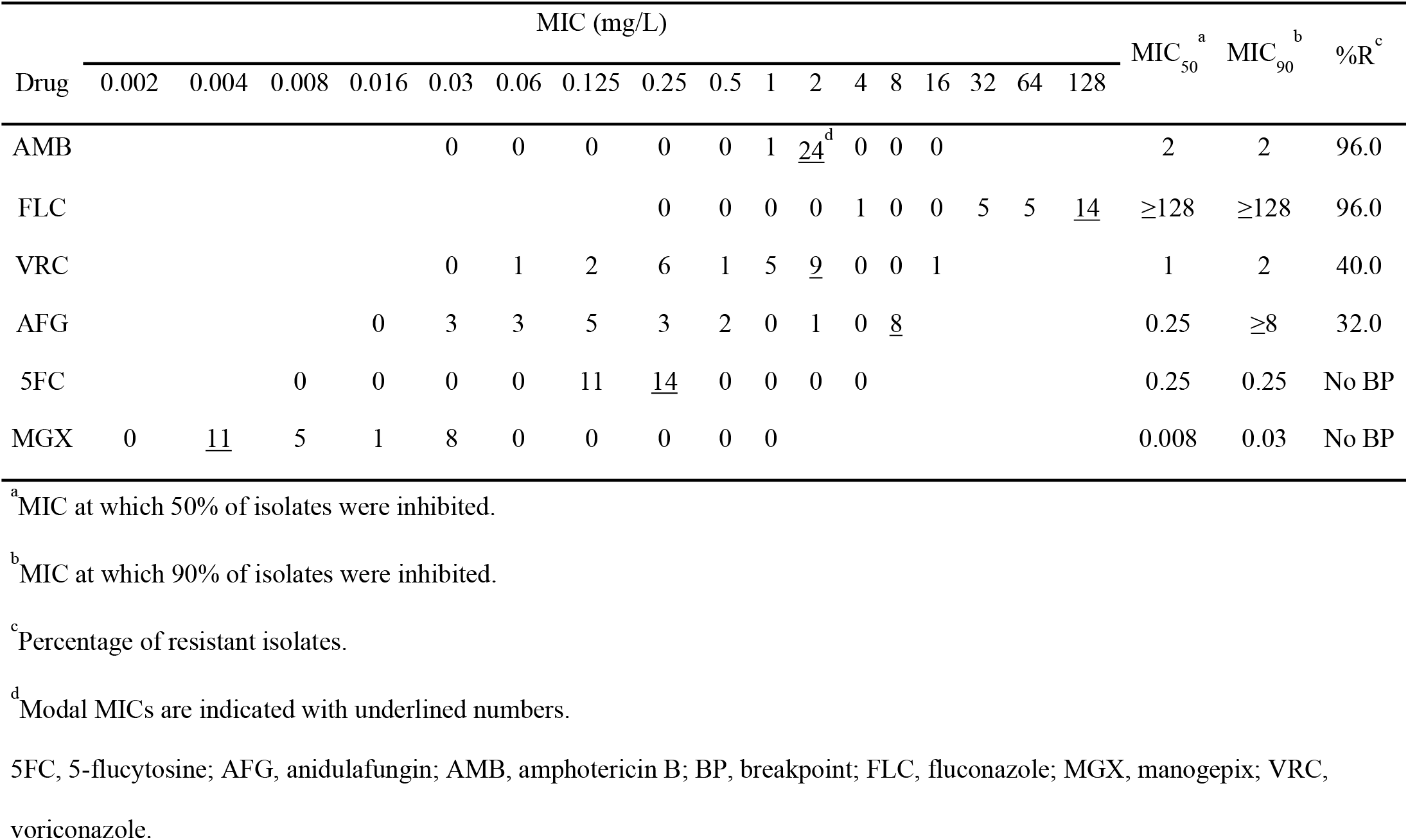
Antifungal MIC distribution for 25 *C. auris* isolates.

### Interaction of antifungal drug combinations against C. auris isolates

Based on their MIC values, 11 *C. auris* isolates with different drug susceptibility profiles were selected to investigate the interactions of anidulafungin, amphotericin B and voriconazole with 5-flucytosine or manogepix. The median FICI values for these combinations, as determined by the checkerboard assays, are presented in Table 3 and Figure 1 (FICI values of separate repeats can be found in Tables S2 and S3). The combinations of anidulafungin with 5-flucytosine or manogepix resulted in synergistic interactions for 10/11 isolates (synergy, 2/11 isolates; partial synergy, 8/11 isolates). Meanwhile the combination of anidulafungin with manogepix led to synergy in all 11 isolates (synergy, 5/11 isolates; partial synergy, 6/11 isolates). These FICI values corresponded to a median (range) decrease in MICs of 2 log_2_-fold (1- to 4 log_2_-fold) for anidulafungin and 2 log_2_-fold (0- to 4 log_2_-fold) for 5-flucytosine, or 3 log_2_-fold (1- to 9 log_2_-fold) for anidulafungin and 2 log_2_-fold (1- to 3 log_2_-fold) for manogepix (Figure 2).

**Table 3.**
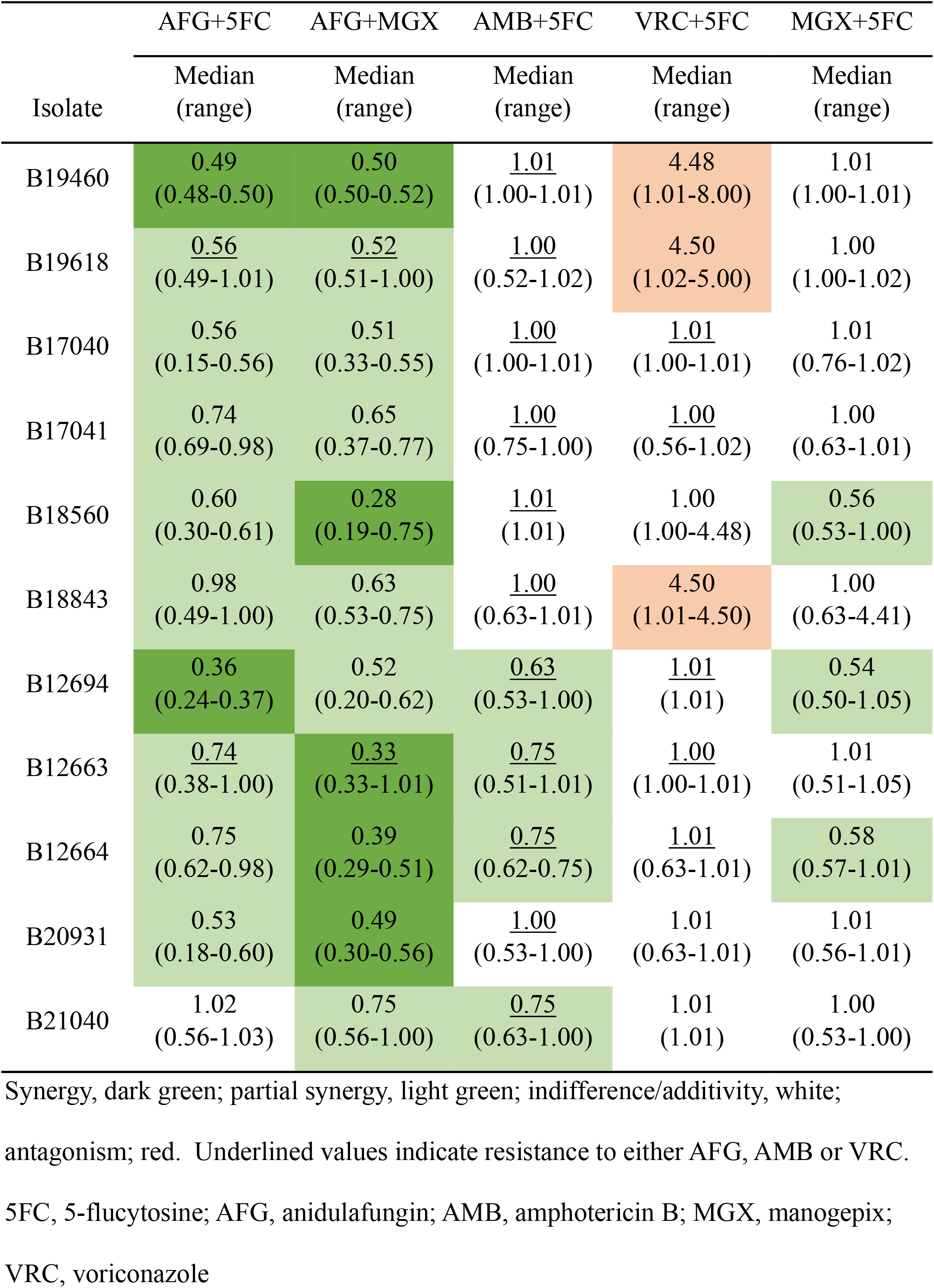
FICI values for 5 antifungal combinations against 11 *C. auris* isolates.

**Figure 1.**
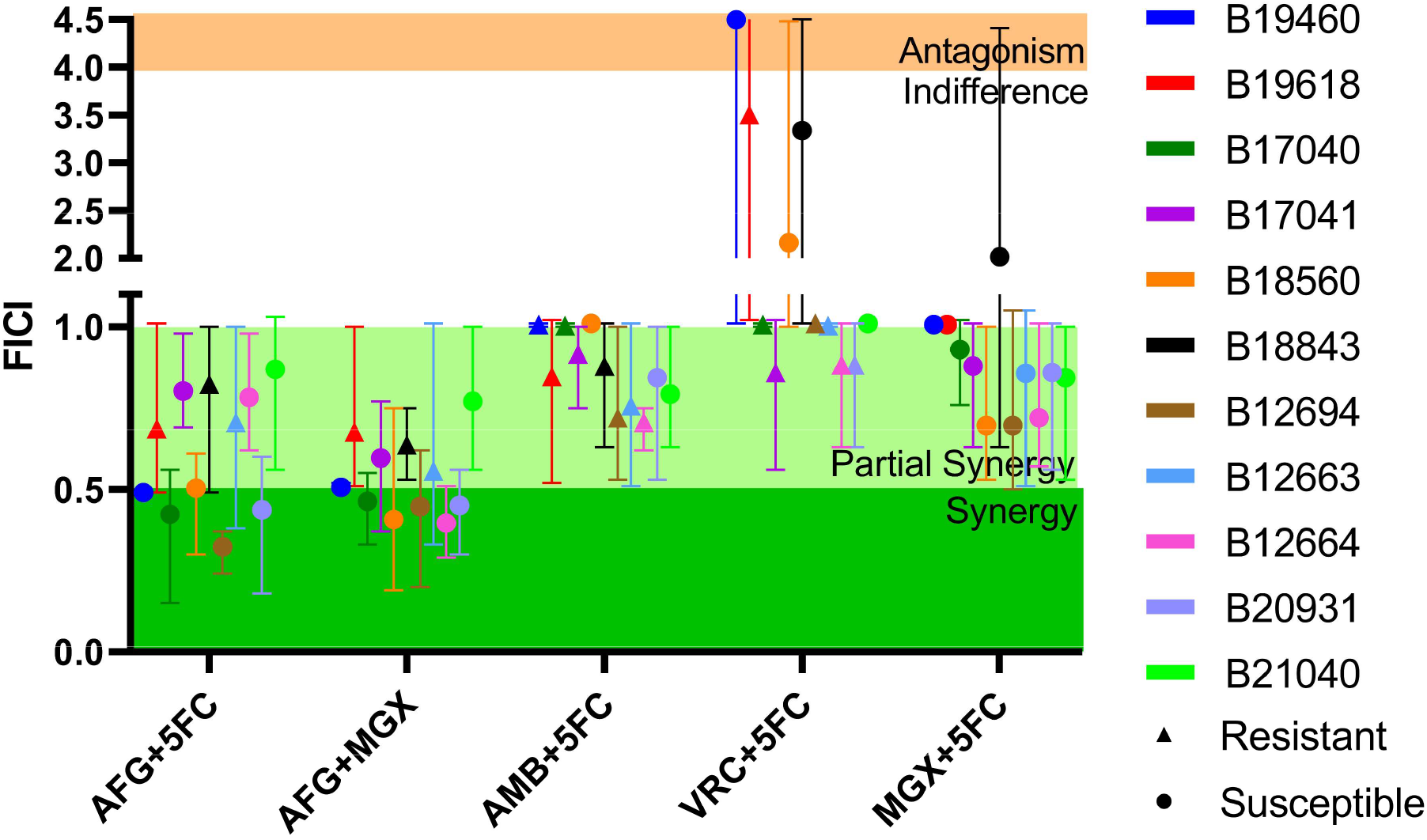
*In vitro* interactions of AFG, MGX, AMB, VRC and 5FC according to the FICI values for 11 *C. auris* isolates. Susceptible (circle) and resistant (triangle) isolates are specified for antifungals with available breakpoints. Minimum FICI values shown in absence of antagonism, otherwise maximum FICI values reported. Drug interaction ranges are indicated by background colour: Synergy, dark green; partial synergy, light green; indifference, white; antagonism, red. Symbols represent median FICI values ± range of three independent experiments. 5FC, 5-flucytosine; AFG, anidulafungin; AMB, amphotericin B; MGX, manogepix; VRC, voriconazole.

**Figure 2.**
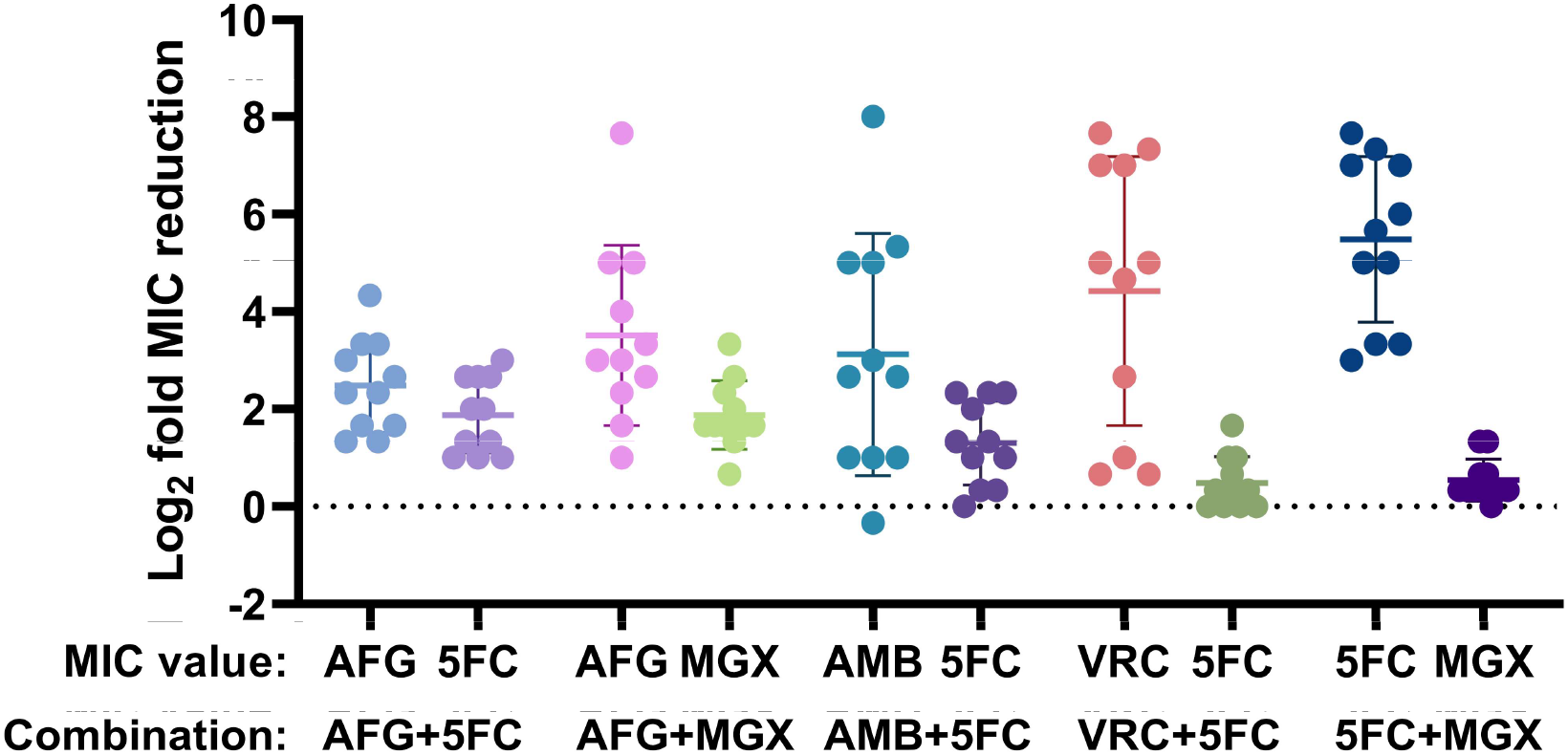
Log_2_ fold reductions in MIC values due to antifungal combinations for 11 *C. auris* isolates. Changes in MIC values for 11 *C. auris* isolates in combinations of anidulafungin, amphotericin B, voriconazole, manogepix and 5-flucytosine compared to the antifungals in monotherapy as determined by checkerboard assays. Symbols represent mean values of three independent experiments. Error bars depict mean ± SD of all isolates per combination. 5FC, 5-flucytosine; AFG, anidulafungin; AMB, amphotericin B; MGX, manogepix; VRC, voriconazole.

The combination of amphotericin B with 5-flucytosine did not show full synergy for any of the tested isolates, though partial synergy was observed in 4/11 isolates (median FICIs 0.63-0.75). The other isolates showed either additive (5/11 isolates) or indifferent (2/11 isolates, median FICIs 1.01) interactions for amphotericin B with 5-flucytosine. For the combination of manogepix and 5-flucytosine, 3/11 isolates displayed partial synergy (median FICIs 0.54-0.58) and 4/11 isolates showed additive or indifferent interactions (median FICIs 1.01). The combination of manogepix and 5-flucytosine led to large reductions in the MICs by median (range) 7 log_2_-fold (1- to 8 log_2_-fold) for 5-flucytosine, while the manogepix MICs were only decreased by median (range) 0 log_2_-fold (0- to 2 log_2_-fold). The drug combination resulting in the least favourable interactions was voriconazole with 5-flucytosine with 3/11 isolates displaying antagonistic interactions (median FICIs 4.48-4.50), and the remaining isolates displaying additive (3/11 isolates) or indifferent (5/11 isolates, median FICIs 1.01) interactions.

Response surface analyses were also used to examine the drug combinations, and an example is shown in Figure 3 for the multidrug-resistant isolate B12663 (see Figures S1-S5 for the other isolates). Consistent with the FICI scores, the synergy maps indicate synergy for the combination of anidulafungin and manogepix (median FICI 0.33) and weak synergy for combinations of 5-flucytosine with anidulafungin (median FICI 0.74) or amphotericin B (median FICI 0.75). In contrast to the FICI calculation, which only focuses on drug concentrations corresponding to MIC values, the response surface analysis permits the examination of drug interactions over a wide range of tested concentrations. This revealed antagonism at the lower end of some concentration ranges that was missed by the FICI approach, highlighting the concentration-dependence of the interactions.

**Figure 3.**
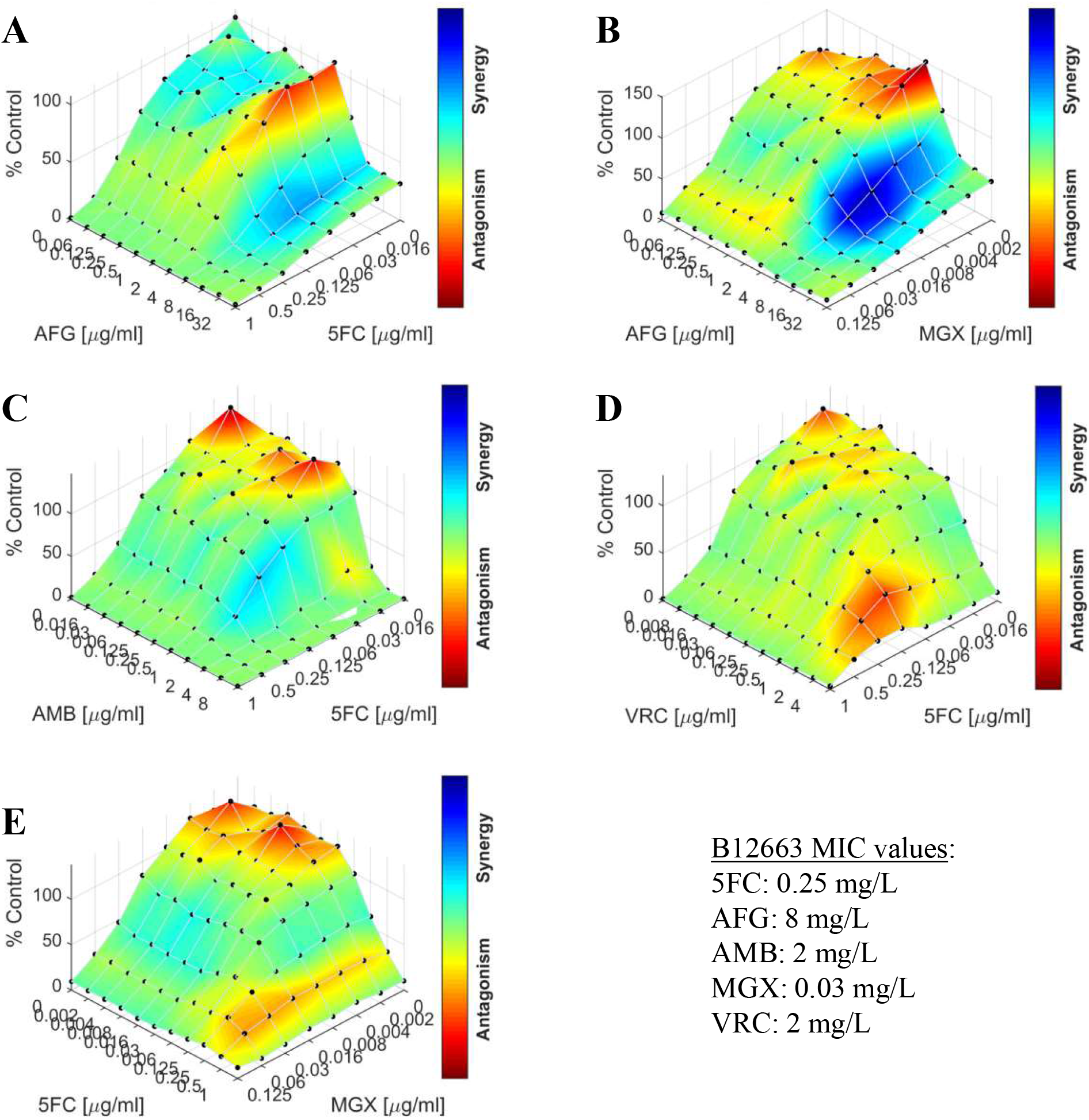
Synergy maps for 5 antifungal combinations against the multidrug-resistant *C. auris* isolate B12663. The interactions of 5-flucytosine with anidulafungin (A), amphotericin B (C) or voriconazole (D) and the interactions of manogepix with anidulafungin (B) or 5-flucytosine (E) were analysed with Combenefit (n=3). The graphs show the growth percentage relative to the drug-free control with the colour scale representing the drug interaction. 5FC, 5-flucytosine; AFG, anidulafungin; AMB, amphotericin B; MGX, manogepix; VRC, voriconazole.

### Real time imaging of anidulafungin combinations against a multidrug-resistant C. auris isolate using microfluidics

A microfluidics imaging approach was employed to further investigate the effects, at a single-cell level, of the two most promising drug combinations: anidulafungin with manogepix, and anidulafungin with 5-flucytosine. This system is less static than the traditional microbroth dilution method as the cells are constantly perfused with fresh medium containing different antifungal drugs. Again, the multidrug-resistant *C. auris* isolate B12663 was chosen for analysis. Both drug combinations showed dramatic effects upon cell growth, markedly reducing the size of colonies compared to the relevant monotherapies and media-only controls (Figure 4A; Movies S1 and S2). Doubling times increased significantly in the presence of the drug combinations compared to the individual antifungals. An increase from 3.19 h (5-flucytosine alone) to 4.90 h (p<0.001) was observed for anidulafungin combined with 5-flucytosine (Figure 4B). Similarly, an increase from 2.75 h (manogepix alone) to 9.50 h (p<0.001) was seen for the anidulafungin-manogepix combination (Figure 4C). These changes in doubling time correspond to 63.5 % (anidulafungin-5-flucytosine) and 96.5 % (anidulafungin-manogepix) decrease in colony area after 24 h compared to 5-flucytosine and manogepix, respectively (data not shown). These findings were again consistent with those of the checkerboard and response surface analysis experiments, in that the combination of anidulafungin and manogepix showed the most potent impacts on cell growth, followed by the combination of anidulafungin plus 5-flucytosine.

**Figure 4.**
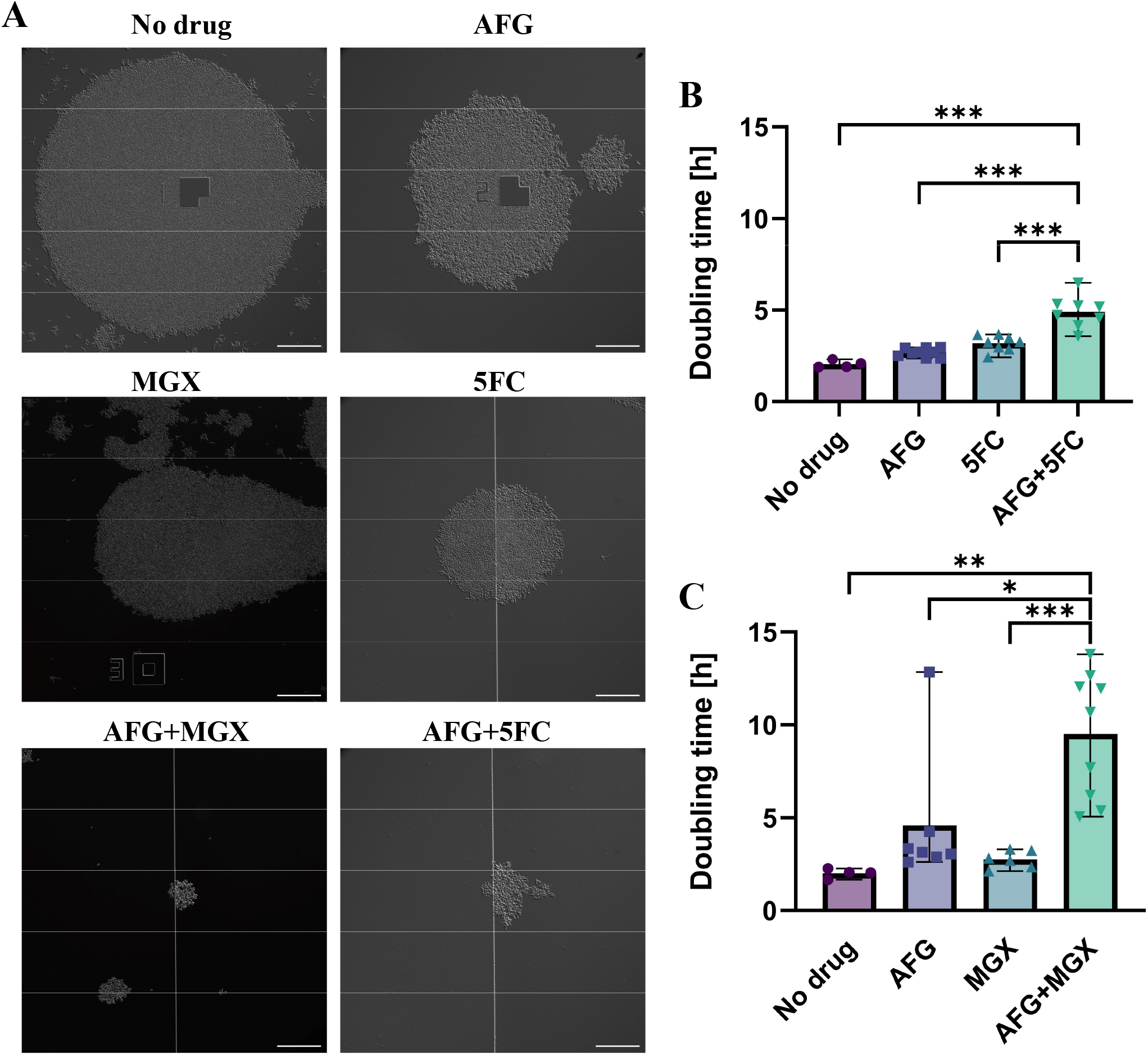
Microfluidics imaging of *C. auris* under antifungal combination exposure. DIC images from two representative experiments (A) and doubling times (B, C) of *C. auris* B12663 cells grown in the presence of RPMI 2%G for 4 h, followed by further RPMI 2%G or treatment with anidulafungin, 5-flucytosine and manogepix alone or in combination at their MICs for 16 h. Doubling times were calculated for several colonies from two independent experiments. Mean ± range. Scale bars: 100 μm. *P≤0.05; **P ≤ 0.01; ***P<0.001 (one-way ANOVA test with Bonferroni’s correction). 5FC, 5-flucytosine; AFG, anidulafungin; MGX, manogepix.

## Discussion

The emergence and global spread of multidrug-resistant *C. auris* strains poses a serious health threat. The high prevalence of antifungal resistance reported for *C. auris* isolates (3, 6–9, 11, 18, 24) was also observed in the isolates used in this study, with the majority of isolates resistant to fluconazole, 40 % resistant to voriconazole and 32 % resistant to anidulafungin. The ability of *C. auris* to develop resistance to all of the available classes of antifungal drug severely limits treatment options.

New antifungal drugs, such as fosmanogepix, are currently in development (reviewed in (30)). *C. auris* currently appears susceptible to the active version of this new class of drugs (manogepix), but there is a high risk of resistance developing following its introduction to the clinic unless precautionary measures are taken. Combination therapies provide a proven strategy that has already been employed in the treatment of viral and bacterial infections to prevent the emergence of resistance to a single drug (37). Additionally, combination therapies have the potential to improve efficacy through additive or synergistic interactions, allowing lower drug doses to be used, thereby reducing dose-related toxicity.

Thus far, nine studies have examined antifungal drug combinations against *C. auris*. The majority of these studies focussed on combinations of azoles with echinocandins (20, 23, 24, 38), while a smaller number have evaluated polyene-echinocandin interactions (21, 22) or combinations with 5-flucytosine (25–27). These studies reported mainly synergistic (including partial synergy) or indifferent interactions, with inter-strain variability observed for some combinations. None of these studies included manogepix. Both manogepix and 5-flucytosine have potent antifungal activity against *C. auris* as shown here and observed by others (39–44). Therefore, we examined interactions of the echinocandin anidulafungin, the azole voriconazole and the polyene amphotericin B with either 5-flucytosine or manogepix using checkerboard assays, response surface analyses and microfluidics imaging.

According to the FICI values and response-surface analyses, the most potent combination (with respect to the number of *C. auris* isolates that displayed synergy) was anidulafungin plus manogepix, followed by the combination of anidulafungin with 5-flucytosine. The high efficacy of these combinations was also confirmed by microfluidics imaging, which revealed dramatic reductions in fungal growth compared to the relevant monotherapies. The interactions between 5-flucytosine with either amphotericin B or manogepix were additive or indifferent for the majority of the isolates, while the combination of voriconazole with 5-flucytosine was indifferent or antagonistic.

Applying our FICI thresholds, Bidaud and co-workers also reported mainly partially synergistic or additive interactions for combinations of amphotericin B, voriconazole or micafungin with 5-flucytosine (25). However, they did not observe the antagonism for the combination of voriconazole with 5-flucytosine that we observed here. Another study reported 100 % growth inhibition of amphotericin B or anidulafungin-resistant *C. auris* isolates for amphotericin B-5-flucytosine combinations (0.25/1 mg/L) or anidulafungin-5-flucytosine combinations (0.008/1 mg/L) (26). Based on our OD_530_ measurements, more than 90 % growth inhibition was also achieved for the majority of susceptible and resistant isolates we analysed, and this growth inhibition could be reached at lower concentrations for some isolates. To the best of our knowledge, antifungal combinations with fosmanogepix/manogepix have not been studied previously against *Candida* species. One recent study compared amphotericin B monotherapy with the combination therapy of fosmanogepix and amphotericin B in invasive mouse infection models of *Aspergillus fumigatus, Rhizopus arrhizus* var. *delemar* and *Fusarium solani* (45). In all three models, mortality and fungal burden were significantly reduced in the mice treated with the combination therapy compared to amphotericin B or fosmanogepix alone (45).

For the majority of combinations and isolates we examined, the interactions were partially synergistic or additive. However, even these interactions could be of interest clinically, as the ultimate goal is to reduce fungal burden with a view to supporting the immune system in clearing the infection. This reduction in fungal growth could be clearly observed in the microfluidics imaging for the combination of anidulafungin with 5-flucytosine, which only displayed a partially synergistic interaction for the imaged isolate in the checkerboard assays. Furthermore, partially synergistic or additive interactions can lead to reductions in the MICs, potentially allowing for a lowering of antifungal doses, thereby reducing toxicity. Reductions in MICs for partially synergistic, additive and indifferent combinations have also been observed by others (20, 24) and Caballero and colleagues reported that additive combinations of isavuconazole-echinocandin combinations against *C. auris* can result in fungistatic effects which were absent for single agents in time-kill assays (23).

It should be noted that the current study employed a relatively small number of isolates, and there was an unequal representation of *C. auris* clades. Additionally, the clustering of amphotericin B MICs around the breakpoint made it difficult to categorise the isolates according to their amphotericin B susceptibility. Hence, other susceptibility testing methods such as the Etest are recommended (17).

In summary, combinations of anidulafungin with manogepix or 5-flucytosine show the highest potential against the tested *C. auris* isolates. Further studies are needed to determine the mechanisms that underlie these drug interactions and to evaluate their efficacy and safety in the murine model and whether these combinations also protect against the development of resistance.

## Acknowledgements

We thank Bethany McCann and Rebecca Inman for helpful discussions and their comments on the manuscript. We thank Joe Sexton, Elizabeth Berkow, Julian Williams, Lynn Kelly, Shawn Lockhart, Anastasia Litvintseva and Tom Chiller at the Centres for Disease Control in Atlanta; Robert Deiss and Sharon Reed from the UC San Diego School of Medicine and Grace Kang from the San Diego County Health and Human Services Agency for sharing and organising the transfer of the clinical *C. auris* isolates used in this study. We also thank Mark Fraser from the UKHSA Mycology Reference Laboratory Bristol for the provision of the quality control strains *C*a*ndida krusei* ATCC 6258 and *Candida parapsilosis* ATCC 22019. This work was supported by the Medical Research Council Centre for Medical Mycology [MR/N006364/2]. EB was also funded by a UK Medical Research Council (MRC) project grant [MR/S001824/1] and Biotechnology and Biological Sciences Research Council (BBSRC) project grant [BB/V017004/1]. AJPB was also funded by an MRC programme grant [MR/M026663/2]. TB was also funded by a Medical Research Foundation Emerging Leaders in Antimicrobial Resistance Award [MRF-160-0009-ELP-BICA-C0802].

